# A feedback-loop between telomerase activity and stemness factors regulates PDAC stem cells

**DOI:** 10.1101/2020.11.02.361931

**Authors:** Karolin Walter, Eva Rodriguez-Aznar, Monica S. Ventura Ferreira, Pierre-Olivier Frappart, Tabea Dittrich, Kanishka Tiwary, Sabine Meessen, Laura Lerma, Nora Daiss, Lucas-Alexander Schulte, Frank Arnold, Valentyn Usachov, Ninel Azoitei, Mert Erkan, Andre Lechel, Tim H. Brümmendorf, Thomas Seufferlein, Alexander Kleger, Enrique Tabarés, Cagatay Günes, Fabian Beier, Bruno Sainz, Patrick C. Hermann

## Abstract

To date, it is still unclear how cancer stem cells (CSCs) regulate their stemness properties, and to what extent they share common features with normal stem cells. Telomerase regulation is a key factor in stem cell maintenance. In this study, we investigate how telomerase regulation affects cancer stem cell biology in pancreatic ductal adenocarcinoma (PDAC), and delineate the mechanisms by which telomerase activity and CSC properties are linked. Using primary patient-derived pancreatic cancer cells, we show that CSCs have higher telomerase activity and longer telomeres than bulk tumor cells. Inhibition of telomerase activity, using genetic *TERT*-knockdown or pharmacological inhibitor (BIBR1532) resulted in CSC marker depletion *in vitro*, and reduced tumorigenicity *in vivo*. Furthermore, we identify a positive feedback loop between stemness factors (KLF4, SOX2, OCT3/4, NANOG) and telomerase, which is essential for the self-renewal of pancreatic CSCs. Disruption the balance between telomerase activity and stemness factors, eliminates CSCs via induction of DNA damage and apoptosis, opening future perspectives to avoid CSC driven therapy resistance and tumor relapse in PDAC patients.

## Introduction

Pancreatic ductal adenocarcinoma (PDAC) is the most frequent and the most lethal form of pancreatic cancer, and is expected to be the second most frequent cause of cancer-related death by 2030 (Rahib *et al*, 2014). Diagnosis of PDAC is frequently delayed due to the absence of early symptoms, pronounced resistance to therapy and early metastatic spread. As a consequence, less than 20% of patients diagnosed with PDAC are eligible for resection (Siegel *et al*, 2015), the only curative treatment option. Despite growing knowledge about PDAC tumor biology, advances in treatment are still scarce, with FOLFIRINOX and gemcitabine + nab-paclitaxel currently representing the most promising combination chemotherapies (Conroy *et al*, 2011; Von Hoff *et al*, 2013). The complexity and heterogeneity of pancreatic cancer remains only partially deciphered, and strategies for developing novel and more effective treatments are urgently needed.

Cancer stem cells (CSCs) have been implicated in a wide variety of tumors. We and others have previously demonstrated their outstanding importance in pancreatic cancer perpetuation, metastasis, and therapy resistance.(Hermann *et al*, 2007; Li *et al*, 2007) While self-renewal is a defining characteristic of CSCs, (Clarke *et al*, 2006) the precise mechanism how CSCs maintain their stemness state remains unclear. Therefore, understanding CSCs at the molecular level might reveal targetable principles that could potentially be exploited therapeutically. Since telomerase activity and telomere regulation have been proposed to play an essential role in tumor cell maintenance and are considered “hallmarks of cancer” by Hanahan and Weinberg, telomerase activity might present such a potential target.

Vertebrate telomeres consist of repetitive TTAGGG DNA sequences. The telomerase complex consists of a catalytic subunit (telomerase reverse transcriptase, TERT) and a telomerase RNA component (TERC). TERC serves as a template for the addition of DNA tandem repeats catalyzed by TERT, which is the rate-limiting factor for telomerase activity. (Nakamura, 1997) Further factors such as the shelterin proteins TERF1 and TERF2 represent important factors for telomerase regulation. (DeLange, 2005) Telomeres protect the ends of the chromosomes from end-to-end fusion and exonucleolytic degradation, preventing genome instability. Importantly, in most human cells telomeric DNA is shortened with each cell division, leading to undetectable telomerase activity in the vast majority of somatic tissues (Wright *et al*, 1996) and accumulation of critically short telomeres, subsequently limiting cellular replicative capacity and ultimately resulting in replicative senescence. (Collado *et al*, 2007)

Telomere length stabilization by the reactivation of telomerase or (much less frequently) through alternative mechanisms of telomere lengthening (ALT) results in elevated telomerase activity in 85-90% of human tumors, highlighting this as the prime mechanism to maintain telomere functionality in cancer. (Shay & Wright, 2019)

Several studies have examined the effects of telomerase inhibition on tumor growth. In cancer cell lines, the potent non-nucleosidic telomerase inhibitor BIBR1532 induces telomere shortening, proliferation arrest and senescence (Damm *et al*, 2001). Currently the effects of the clinical grade telomerase inhibitor GRN163L (Imetelstat) are being tested in several tumors, such as breast and lung cancer, myeloma. However, the effects of telomerase inhibition on CSCs have yet to be elucidated.

In the present study we investigate the role of telomere regulation and telomerase inhibition in the maintenance of pancreatic CSCs. We demonstrate that CSCs isolated from patient-derived xenografts (PDXs) present higher telomerase activity, resulting in significantly longer telomeres compared to bulk tumor cells. Intriguingly, the lengthening of telomeres is inextricably linked to increased expression of the pluripotency / stemness factors OCT3/4, SOX2, NANOG and KLF4, and is jointly regulated in a previously undescribed positive feedback loop which is necessary for these cells to escape senescence, and regulates their stemness properties. Furthermore, pharmacological inhibition of telomerase using BIBR1532 as well as genetic knock-down of TERT greatly decreased the CSC frequency of patient-derived PDAC cells *in vitro* and *in vivo* by CSC-specific induction of DNA damage and apoptosis.

## Results

### Telomerase activity and telomere length in pancreatic CSCs vs bulk tumor cells

In a first screen we tested several primary PDAC cell lines for their average telomere length. Indeed, we observed that all tested samples showed very short telomeres (< 5th age percentile) (**Fig. 1A**). In order to evaluate telomerase activity specifically in CSCs, we compared the expression levels of *TERT* and *TERF1* in CSCs versus bulk tumor cells in three human PDX-derived primary PDAC cell lines (Panc215, Panc185, Panc354). Using well-established CSC enrichment methods (i.e. CD133 ,or Aldefluor fluorescence activated cell sorting (FACS), and sphere culture **(Fig. 1B)**), we found increased expression of *TERT* and/or *TERF1* in CD133+ cells (**Fig. 1C**), Aldefluor+ cells (**Fig. 1D**) and in sphere cultures (**Fig. 1E**) compared to the negative population or adherent cells, respectively. In immunofluorescence staining, significantly more TERT-expressing cells were detectable in the CD133+ CSC subpopulation as compared to CD133− cells (**Fig. 1F**). Next, we measured telomerase activity as the functionally relevant readout. Using gel-based as well as PCR-based TRAP (telomeric repeat amplification protocol) assays, significantly higher telomerase activity was detected in the CSC fraction as compared to the respective control cells (**Fig. 1G-I**). Since the key function of telomerase is telomere elongation, we next performed Q-FISH-analysis to determine telomere length in CD133+ (**Fig. 1J**) or ALDH+ (**Supp Fig. S1A**) CSCs and in the respective marker-negative control populations (Panc215: CD133+ 66.2 a.u. vs CD133− 31.7 a.u.; Panc185: CD133+ 40.8 a.u. vs CD133− 31.5 a.u.; Panc354: CD133+ 41.6 a.u. vs CD133− 35.4 a.u.; and Panc286: ALDH+ 11.2556 a.u. vs ALDH-9.1304 a.u.). Indeed, we observed significantly longer telomeres in the investigated CSC populations. These results confirm increased telomerase expression and activity in pancreatic CSCs as compared to non-CSCs.

**Figure 1.**
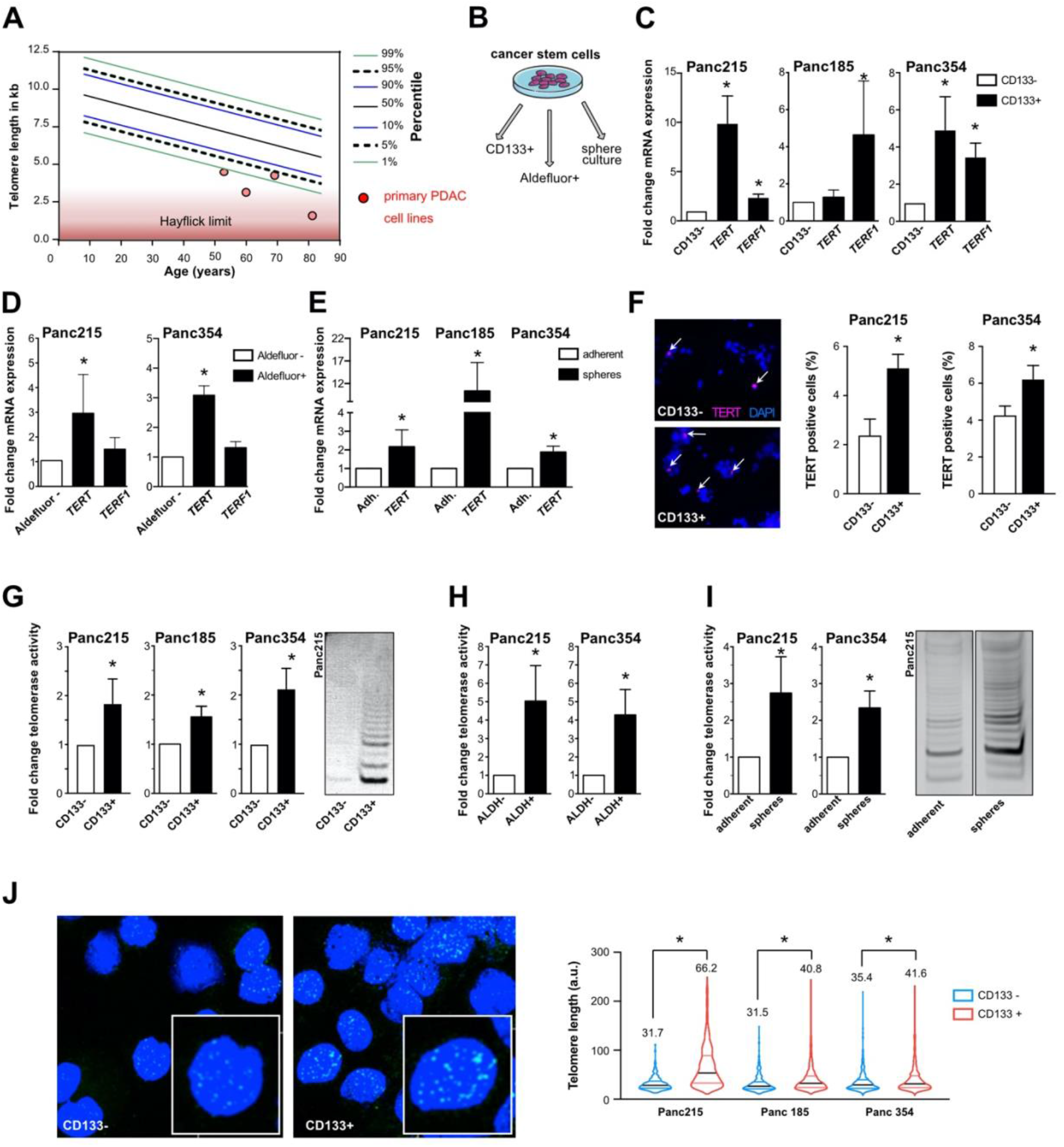
Telomerase activity stabilizes telomere length in pancreatic CSCs. A Telomere length in primary PDAC cells (red dot) with respect to patient age measured by FlowFISH. B Schematic illustration showing the CSC enrichment methods for CD133 and Aldefluor via FACS or in sphere culture. C, D, E RT-qPCR analysis of *TERT* and *TERF1* mRNA levels in primary pancreatic cancer stem cells enriched and selected by FACS for CD133 (C), or Aldefluor (D), and cultured as spheres (E) (CSCs) vs corresponding control (non CSCs) (n = at least 3 independent experiments). F Immunofluorescence staining and quantification for TERT (red) in CD133− and CD133+ FACSorted cells (n = 5 independent experiments). Cells were counterstained with DAPI (nuclear marker, blue). G, H, I Telomerase activity measurement in CD133 (G) or Aldefluor (H) negative vs positive cells and in adherent vs sphere cell-cultures (I) (n = 3 independent FACSortings and sphere culture experiments). J Representative pictures of Q-FISH with telomeres (green) and DAPI (blue) and violin plot showing telomere length analysis in CD133 negative (non CSCs) and positive cells (CSCs). The mean is depicted in numbers and as black line. n ≥ 3 from independent FACSortings with each >150 measurements per group. Data information: In (C-J), data are presented as mean ± SEM. *P ≤ 0.05 (Mann-Whitney-U test). CSC, cancer stem cells; FACS, fluorescence activated cell sorting; FlowFISH, Flow fluorescence in-situ hybridization; PDAC, pancreatic ductal adenocarcinoma; Q-FISH, quantitative fluorescent *in situ* hybridization; RT-qPCR, reverse transcriptase polymerase chain reaction.

### A positive feedback loop between stemness factors and telomerase maintains CSC phenotype

Since we observed a significant increase in the regulation and activity of telomerase in CSCs, we next investigated whether increased telomerase activity is associated with the expression of genes and markers of stemness and pluripotency. We observed increased expression of *OCT3/4*, *NANOG*, and *LGR5* in the CSC population (**Fig. 2A**) as compared to the CD133− control population. Similarly, expression levels of these genes and of *SOX2* were significantly elevated in Aldefluor-positive cells (**Fig. 2B**) compared to the negative cells, demonstrating increased expression of most stemness-associated genes in CSCs with some variation depending on the utilized identification strategy. To overcome CSC marker-dependent variation, we next infected Panc354 primary cell cultures with a Nanog-YNL reporter system. (Takai A., 2015) YNL+ cells showed increased expression of *NANOG*, but also of *TERT* and other stemness-associated genes (**Fig. 2C**). Furthermore, YNL+ cells showed significantly higher telomerase activity (**Fig. 2D**). While the amount of CD133+ cells was unchanged (**Fig. 2E**), sphere forming capacity was significantly increased (**Fig. 2F**) in the YNL+ cells. To confirm higher telomerase activity in CSCs, we used a recombinant pseudorabies virus (PRV) that we previously designed and demonstrated to replicate only in cells with high telomerase activity (PRV-TER). (Lerma *et al*, 2016) Adherent cultures of human pancreatic ductal epithelial (HPDE) cells and Panc185 cells were infected with two parental control viruses (PRV-NIA3 and vBecker2) and with PRV-TER. While both parental viruses efficiently replicated in HPDE and Panc185 cultures, PRV-TER was unable to produce *de novo* virions in either culture (**Fig. 2G**). In contrast, when CSC-enriched sphere cultures were infected, all three viruses efficiently replicated and produced *de novo* virions, confirming that CSC-enriched sphere cultures express significantly higher telomerase activity, allowing for PRV-TER to efficiently replicate in these cultures.

**Figure 2.**
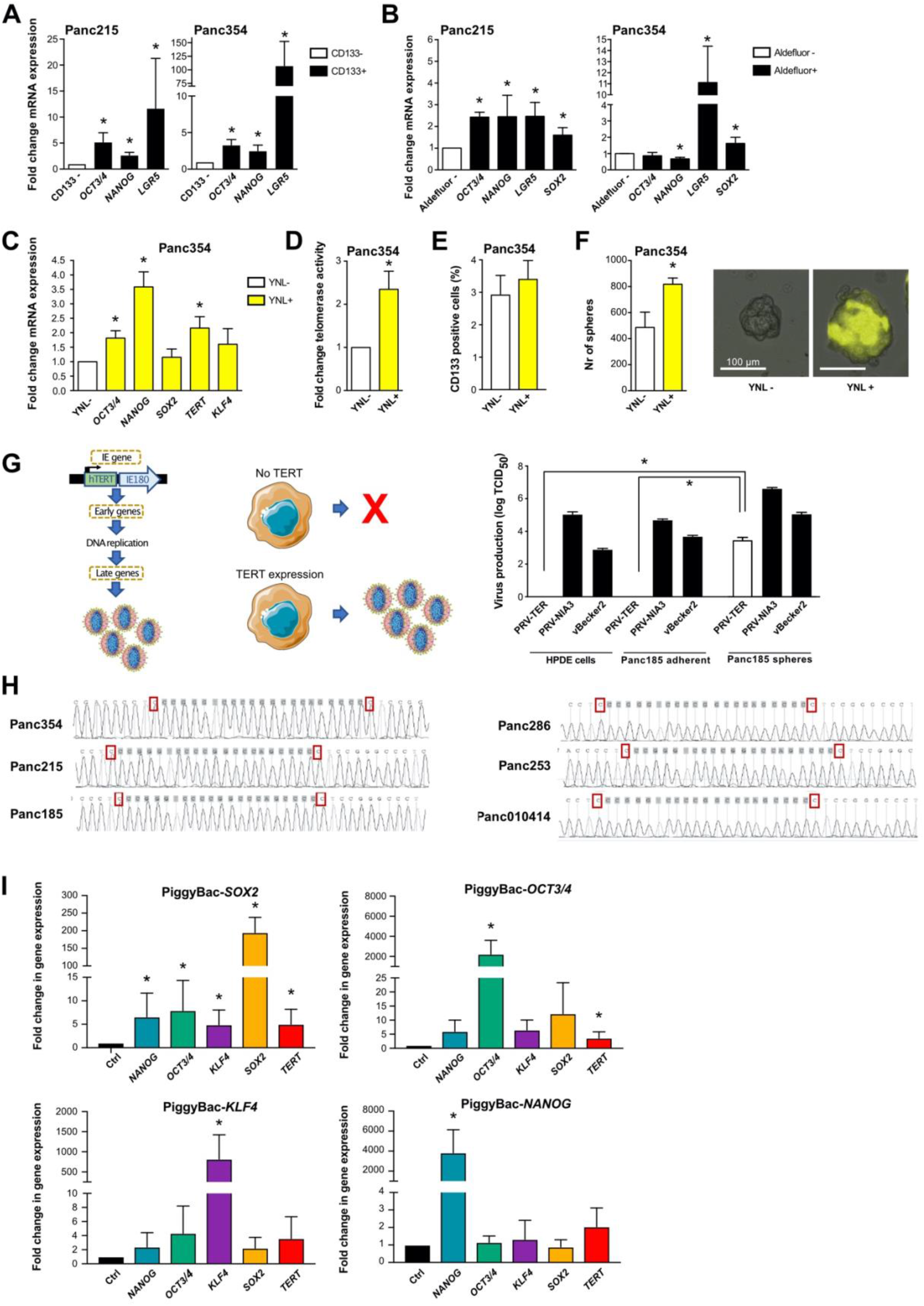
Interplay between pluripotency / stemness factors and telomerase activity in pancreatic CSCs. A, B RT-qPCR analysis of pluripotency / stemness-associated genes in primary pancreatic CSCs enriched by CD133 expression (A) or Aldefluor activity (B) vs CD133− and Aldefluor-non-CSCs (n = at least 3 independent FACSortings). C, D, E, F Quantitative RT-PCR (C), telomerase activity (D), flow cytometry analysis for CD133 (E) and sphere formation capability (F), compared in NANOG negative cells (white) vs NANOG positive cells (yellow) using a NANOG-YNL reporter. G Schematic illustration and quantification on the production of virions in parental pseudorabies viruses (PRV-NIA3 and vBecker) compared to telomerase activity-dependent virus production (PRV-TER) in adherent HPDE and Panc185 cells as compared to Panc185 spheres. H TERT promoter mutation analysis of C250T and C228T sites in 6 primary pancreatic cancer cells (Panc215, Panc354, Panc185, Panc286, Panc253, and Panc010414). I Gene expression levels of stemness / pluripotency genes and *TERT* in HEK293T cells after inducing expression of SOX2, OCT3/4, KLF4 and NANOG, compared to the empty vector control (n = 4 independent experiments). Data information: In (A – G and I), data are represented as mean ± SEM. *P ≤ 0.05 (Mann-Whitney-U test). YNL, yellow Nano-latern reporter system.

Finally, we explored the mechanism(s) underlying increased telomerase regulation in CSCs. Direct promoter activation can lead to increased *TERT* activity in tumor cells, and two common promoter mutations (−124 and −146 bp upstream of the *TERT* start site) have been associated with increased telomerase activity. (Vinagre *et al*, 2013) These mutations are frequent in some cancers, but rarely occur in gastrointestinal cancers. Indeed, promoter sequencing of all tested PDAC cells revealed wildtype sequences (6/6 primary cell lines, 5/5 established cell lines, **Fig. 2H** and **Supp Fig. S1B**).

We next generated PiggyBac expression vector constructs that specifically express *SOX2, OCT3/4, KLF4* or *NANOG* to determine how these stemness factors modulate *TERT* expression. Supporting this approach, Hsieh et al. showed that PARP1 recruits KLF4 to activate telomerase expression in embryonic stem cells. (Hsieh *et al*, 2017) Interestingly, the expression of *OCT3*/4 and *SOX2* significantly upregulated *TERT* expression and a similar trend was observed for *NANOG* and *KLF4* (**Fig. 2I**). Furthermore, overexpression of any of these factors resulted in concomitant upregulation of the other factors as well, indicating an intricate cross-regulation of these individual stemness factors. Taken together, these data show that telomerase activity is increased in CSCs and is regulated by a feedback loop between *TERT* and key pluripotency-associated factors.

### Telomerase inhibition causes DNA damage and apoptosis in CSCs

Based on the observed link between telomerase activity / telomere length and a CSC phenotype, we next evaluated the effects of telomerase inhibition on the CSC population. For this purpose, we treated primary pancreatic cancer cells with the small molecule telomerase inhibitor BIBR1532 and selected 80μM as an effective sub-IC_50_ concentration (**Supp Fig. S1C**). BIBR1523 treatment was performed for 3d or 7d, respectively, and after 7d of treatment telomerase activity was effectively decreased (**Fig. 3A**). Functionally, treatment with BIBR1523 significantly reduced telomere length (**Fig. 3B**) already after 3d. Interestingly, the treatment effect was potentiated with longer treatment in the CD133+ CSC population. However, in the CD133− bulk tumor cell population telomere length increased again after 7d of treatment. This is most likely due to early elimination of CD133− cells with critically short telomeres and the inability of CD133− cells to upregulated telomerase, leading to a relative enrichment in CD133− cells with longer telomeres. Based on these results, treatment with BIBR1523 was performed for 7d in all further experiments.

**Figure 3.**
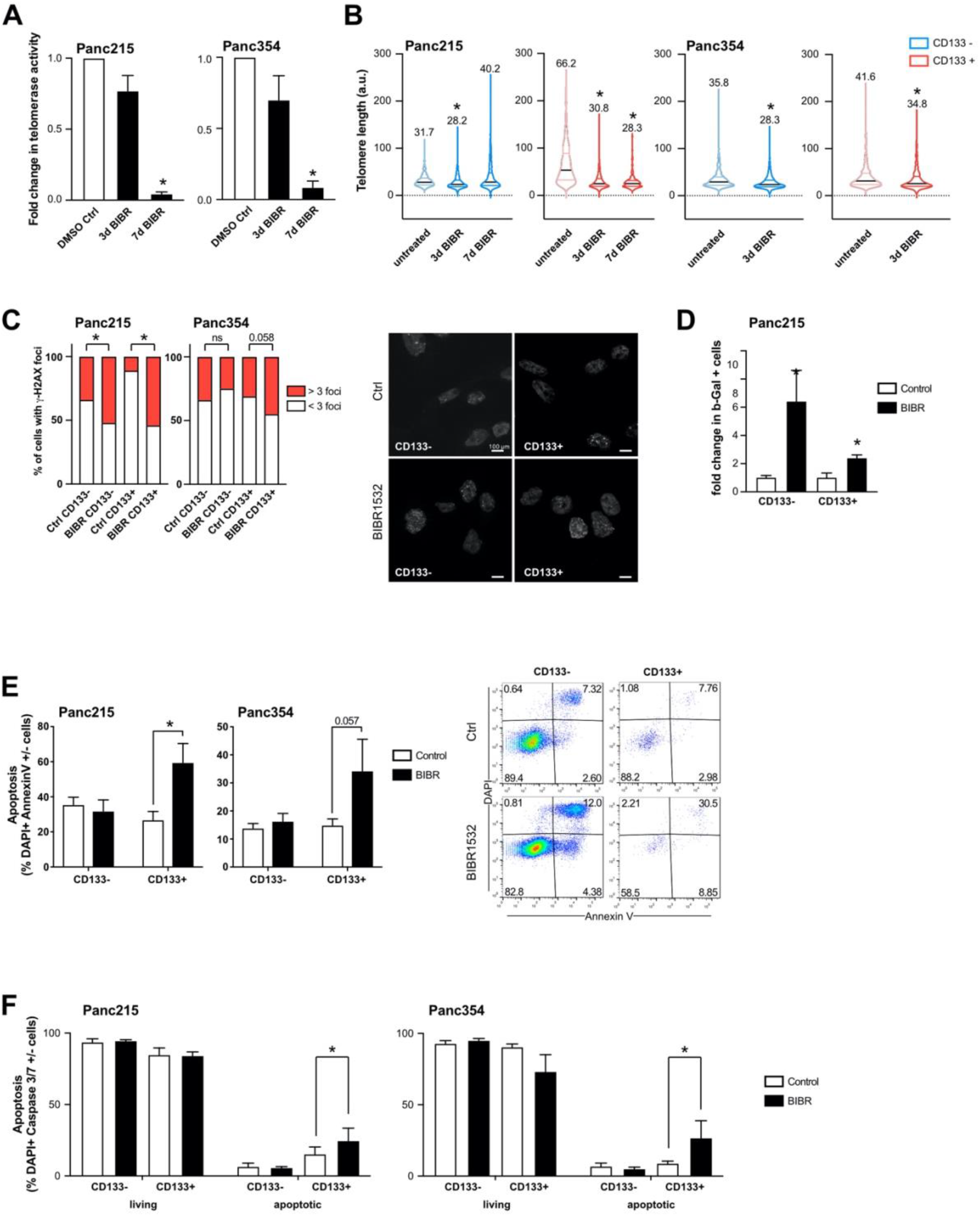
Targeting telomerase activity with a small molecule inhibitor (BIBR1532) A, B The effects of BIBR1532 treatment on telomerase activity (A), and telomere length (B) in CD133− cells (blue) and in the CD133+ CSC population (red) were quantified. For illustrative purposes, the telomere length measurements of Fig1H were re-used here (indicated by transparent color). The mean is depicted in numbers and as black line, >150 measurements per group. C Quantification of gH2AX foci (>50 cells per group) in CD133− and CD133+ cells after 7 days vehicle or BIBR1532 treatment. Quantification and representative pictures of immunofluorescence staining of gH2AX foci are provided. D Senescent cells quantified by flow cytometry staining for SA-beta-Gal in CD133− (non CSCs) and CD133+ (CSCs) after BIBR1532 or solvent treatment (7 days). E, F Apoptosis quantified by flow cytometry using double staining for CD133 and AnnexinV or (E) Caspase3/7 (F). Data information: In (A-F), data are represented as mean ± SEM. n = 3 independent experiments. *P ≤ 0.05 (Mann-Whitney-U test).

To determine the biological consequence of telomerase inhibition, we quantified DNA damage by γ-H2AX staining, as telomerase inhibition can induce DNA damage, cell cycle arrest and apoptosis in other systems. (Celeghin *et al*, 2016) Treatment with BIBR1532 increased γ-H2AX foci generation much more strongly in CD133+ CSCs than in bulk tumor cells, indicating a preferential induction of DNA damage after telomerase inhibition in CSCs (**Fig. 3C**). Accumulation of DNA double strand breaks (DSBs) and the activation of DNA damage response are considered contributing factors to cellular senescence;(Collado *et al.*, 2007) Therefore, we next measured senescence using flow cytometry to detect ß-galactosidase after BIBR1532 treatment and observed increased senescence in both CD133− and CD133+ cells, excluding preferential induction senescence in CSCs as response to BIBR1532 treatment (**Fig. 3D**).

Since DSBs can also induce apoptosis, we next measured early apoptotic cells by AnnexinV staining and observed significantly more apoptotic cells in the CD133+ CSC population upon BIBR1532 treatment, while CD133− cells were unaffected (**Fig. 3E**). Furthermore, we performed Caspase 3/7 staining to determine whether BIBR1532-induced DNA damage leads to the recruitment of apoptosis effector kinases. In line with the AnnexinV staining, BIBR1532 treatment resulted in increased Caspase 3/7 staining in the CD133+ CSC population, while no differences were observed in CD133− cells (**Fig. 3F**).

### CSCs are depleted upon telomerase inhibition

Since BIBR1532 treatment appeared to have CSC-specific effects, we next evaluated the effect of telomerase inhibition on CSC properties. As expected, BIBR1532 treatment strongly reduced the expression of *TERT*, along with established stemness / pluripotency-associated genes (**Fig. 4A**), and significantly decreased the percentage of CSCs as measured by expression of CD133 or Aldefluor (**Fig. 4B, Supp Fig. S1D, S1E**) or sphere formation capacity (**Fig. 4C**). Most importantly, BIBR1532 treatment strongly reduced the number of tumor-initiating cells in (extreme limiting dilution (ELDA) tumorigenicity assays in nude mice. But also the tumor growth of the 100.000 injected cells after BIBR1532 treatment was significantly reduced compared to the tumor growth of solvent treated cells (**Fig. 4D**). These data clearly indicate that telomerase inhibition by BIBR1532 strongly affects the functional capacity of the CSC population.

**Figure 4.**
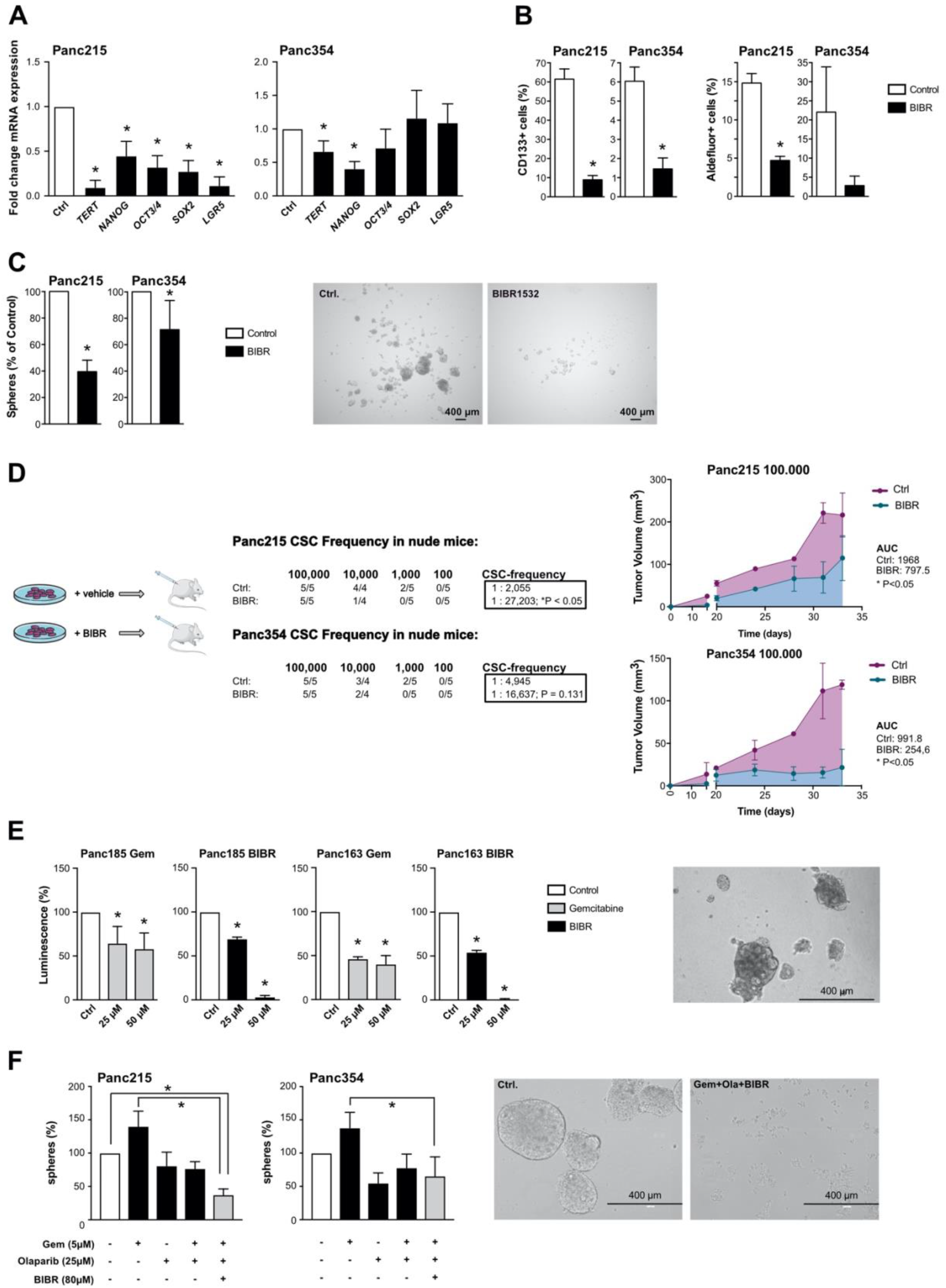
Telomerase inhibition as treatment strategy for pancreatic cancer (stem) cells. A Effects of the small molecule telomerase inhibitor BIBR1532 on the expression of *TERT* and pluripotency-associated genes as measured by RT-qPCR (n = 4 independent experiments). B Flow cytometry analyses for CD133 or Aldefluor after BIBR1532 or solvent treatment (n = 3 independent experiments). C Quantification and representative pictures of spheres after BIBR1532 treatment (n = 3 independent experiments). D Schematic of BIBR1532 pre-treatment and *in vivo* experiment, number of tumorigenic cells within the whole population shown as cancer stem cell (CSC) frequencies as determined by extreme limiting dilution assays (ELDA) in nude mice, and tumor volume measured after injection of 100.000 BIBR1532 or solvent treated Panc215 and Panc354 cells (n ≥ 4 mice for each group). E Viability of PDX-derived organoids treated with gemcitabine and BIBR1532 at the indicated concentrations. F Quantification and representative pictures of sphere formation assays after single agent or combination treatment with gemcitabine, Olaparib, and BIBR1532 (n = 5 independent experiments). Data information: In (A-C; E-F) data are represented as mean ± SEM. *P ≤ 0.05 (Mann-Whitney-U test). In (G) data are represented as mean ± SEM. *P ≤ 0.05 (Area under the curve).

We have previously shown that chemotherapy enriches for CSCs, and have demonstrated the efficacy of combining stem cell inhibitors, stroma-depleting agents and chemotherapy to significantly improve survival of mice *in vivo*.(among others:(Lonardo *et al*, 2011; Mueller *et al*, 2009)) In organoid cultures generated from primary pancreatic cancer cells we observed a significant decrease in luminescence (i.e. viability) upon treatment with increasing doses of gemcitabine. However, gemcitabine did not abrogate organoid formation, which was only achieved using BIBR1532 treatment (**Fig. 4E**). Since the induction of DNA damage by telomerase inhibition did not entirely abrogate CSC function (see **Fig. 3**), we used a combination of chemotherapy, BIBR1532 and the PARP inhibitor Olaparib to target CSCs more successfully. Indeed, evaluating sphere formation as surrogate marker for CSC activity, the combination therapy had significantly stronger effects than chemotherapy alone (**Fig. 4F**).

### Telomerase inhibition as novel treatment strategy for pancreatic cancer

In order to ensure that the effects of BIBR1532 are specific to telomerase inhibition, we proceeded to corroborate our main findings with shRNA-mediated *TERT*-knockdown (*TERT*-KD). As expected, genetic knockdown of *TERT* using two different shRNAs suppressed the expression of *TERT* (**Fig. 5A**), and strongly suppressed telomerase activity (**Fig. 5B**); due to its stronger effects, shRNA 340160 was used for all subsequent experiments. *TERT*-KD suppressed the expression of stemness-associated genes (**Fig. 5C**). The reduction in stemness translated into depletion of CD133+ CSCs (**Fig. 5D**) and reduced sphere formation activity (**Fig. 5E**). Similar to the effects of BIBR1532 (**Fig. 3F&G**), the *TERT*-KD significantly increased apoptosis in CD133+ cells but not in CD133− cells (**Fig. 5F**). Most importantly, the *TERT*-KD strongly reduced TIC frequency in ELDA xenografting assays (**Fig. 5G, Suppl. Fig 1F**). These results closely resemble those achieved with BIBR1532, confirming that the effects observed above are indeed telomerase-specific.

**Figure 5.**
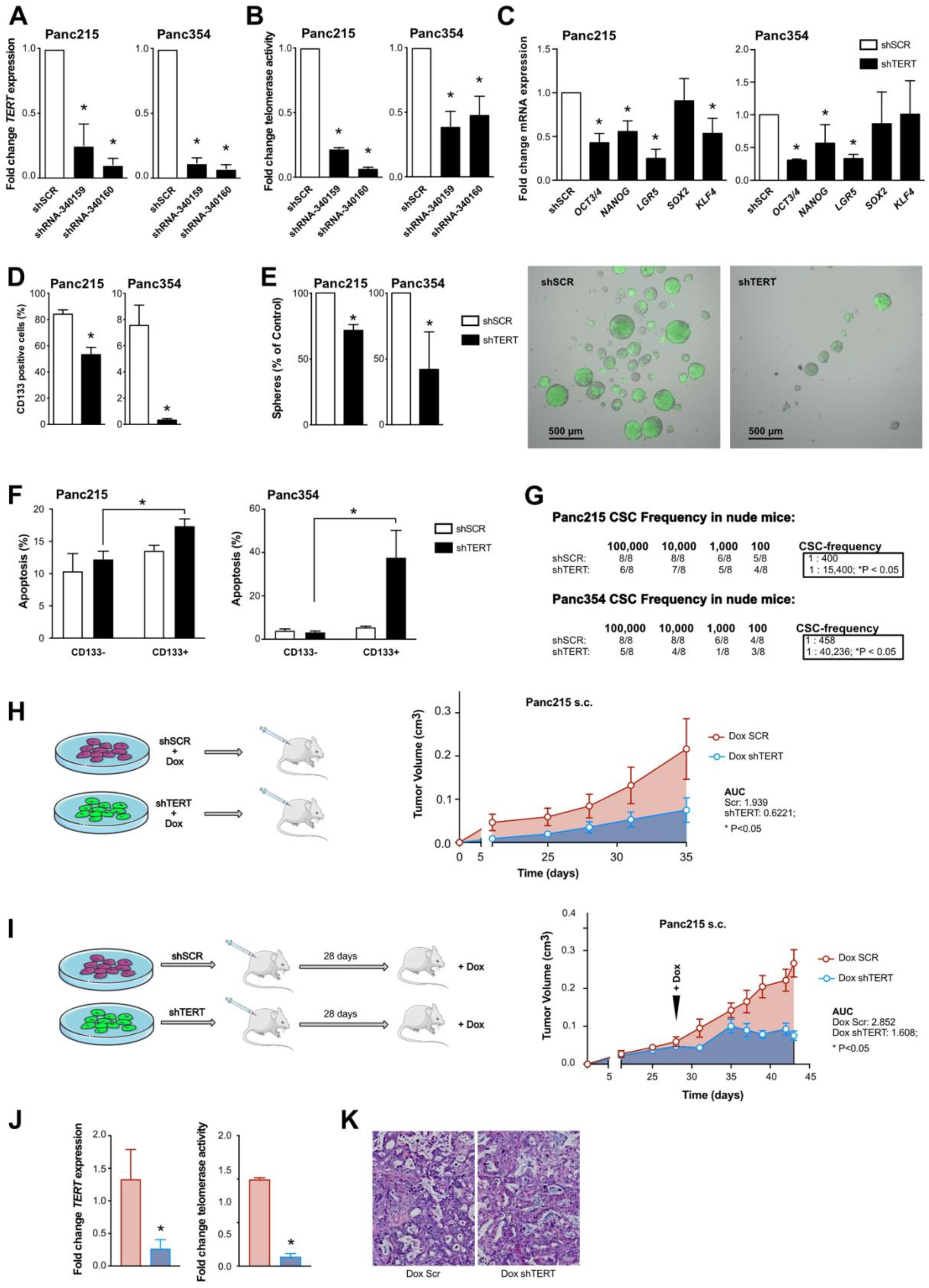
Knockdown of TERT diminishes pancreatic cancer stem cells. A, B Gene expression of *TERT* (A) and telomerase activity (B) in Panc215 and Panc354 cells transduced with shRNA-340159 and shRNA-340160 compared to scrambled (shSCR) control. C Expression levels of stemness / pluripotency-associated genes using shRNA-340160 (shTERT) for *TERT* mediated knockdown in Panc215 and Panc354 cells. D, E Flow cytometry analyses for CD133 (D) and sphere formation (E) upon *TERT* knock-down (KD) compared to scrambled (shSCR) control. F Quantification of apoptosis in shSCR and shTERT transduced cells (Panc215 and Panc354) using flow cytometry analysis for AnnexinV (n = at least 3 independent experiments). G *In vivo* tumor-initiating potential with CSC frequencies (number of tumorigenic cells within the whole population) as determined by ELDA in nude mice after *TERT*-KD compared to scrambled control (n = 8 animals). H Schematic illustration of *in vivo* experiment with cells carrying an inducible *TERT*-KD (Dox shTERT) or scrambled (Dox SCR) control construct. Cells were treated with doxycycline (DOX) 7 days before s.c. xenografting in nude mice. Graph shows time-dependent growth of subcutaneously engrafted tumors (n = 4 mice per group). I Schematic illustration of *in vivo* experiment and visual representation of time-dependent tumor growth of subcutaneously (s.c.) engrafted tumors arising from *TERT*-KD and SCR cells, over the course of doxycycline treatment 28 days after s.c. xenografting in nude mice (n = 6 mice per group). Tumor growth is depicted until the first control mice had to be sacrificed. J Gene expression of *TERT* and telomerase activity compared in *TERT*-KD and SCR tumors (induced with DOX at day 28) after sacrificing mice 43 days after xenografting. K representative H&E staining of at day 43 explanted *TERT*-KD and scrambled tumors (induced with DOX at day 28). Data information: In (A-G and J) data are represented as mean ± SEM. *P ≤ 0.05 (Mann-Whitney-U test). In (H and I) data are represented as mean ± SEM. *P ≤ 0.05 (Area under the curve). AUC, area under the curve; ELDA, extreme limiting dilution assay; DOX, doxycycline; KD, knock-down; s.c.; subcutaneous; shSCR, scrambled control; shTERT, shRNA-340160.

In order to model the effects of telomerase inhibition as an applied treatment *in vivo*, we performed subcutaneous xenografting of Panc215 cells carrying an inducible shTERT cassette. Induction of the *TERT*-KD before injection with doxycycline resulted in significantly smaller tumors compared to scrambled control (**Fig. 5H**). In a preclinical trial we xenografted inducible *TERT*-KD Panc215 cells and induced the knockdown 28 days after implantation, when tumors were firmly established. While dox-treated scrambled cells showed continuous tumor growth leading to sacrifice of the mice, we observed disease stabilization after *TERT*-KD (**Fig. 5I**). The residual tumors showed downregulation of *hTERT* as well as decreased telomerase activity (**Fig. 5J**). Histological analysis of the explanted tumors showed no gross differences between the two treatment groups with regard to tumor morphology, cellularity or stroma composition (**Fig. 5K**). Altogether, these data indicate that TERT/telomerase inhibition depletes CSCs, resulting in disease stabilization in PDAC.

## Discussion

We and others have previously identified CSCs in pancreatic cancer, and have demonstrated that they play a key role in the propagation, chemoresistance and metastasis (Hermann *et al.*, 2007; Li *et al.*, 2007). PDAC remains a disease that is terribly difficult to treat. Thus, eliminating CSCs as a continuous source of tumor growth, resistance and relapse is of utmost importance for improved clinical outcome. While functional differences in CSCs are continuously being discovered, the underlying mechanisms for their acquisition of stemness features are much less clear.

CSCs have been isolated based on different markers and cellular features. We have previously established CD133 expression and functional enrichment via sphere formation as reliable CSC markers in PDAC, (Hermann *et al.*, 2007) and together with Aldefluor activity these are most widely used. In addition to using these markers, we evaluated the expression levels of pluripotency-associated genes such as *NANOG, OCT3/4, KLF4*, *SOX2* and others to overcome the limitations of the markers mentioned above, and additionally used a NANOG reporter system to further demonstrate stemness in the isolated CSCs.

Unlimited proliferation requiring telomere maintenance is a “hallmark of cancer” as indicated by Hanahan & Weinberg, and selective telomere maintenance or even elongation via telomerase activity is an essential feature of stem cells. However, available data on telomerase activity and its regulation in CSCs is scarce.

Increased telomere length has been reported in several tumor entities, such as prostate cancer and glioblastoma. (Beier *et al*, 2011; Marian *et al*, 2010) In contrast to these studies and our data, Joseph et al. surprisingly observed no such differences in breast and pancreatic cancer cells. (Joseph *et al*, 2010) This might be explained by the use of only one established cell line versus the panel of primary cells used in our study, as well as by inter-tumoral heterogeneity. Interestingly, Joseph et al. did observe reduced tumor engraftment in nude mice under treatment with the telomerase inhibitor Imetelstat, indicating an effect on a CSC population. While a telomerase-independent mechanism of regulation was suggested by Joseph et al., and the use of high doses of BIBR1532 has also been shown to induce cytotoxicity irrespective of telomerase activity, (El-Daly *et al*, 2005) our data clearly demonstrate the dependency of CSCs on intact telomere regulation. Specifically, we demonstrate that pharmacological inhibition or shRNA-mediated *TERT*-KD result in CSC depletion as evidenced by reduced expression of stemness genes and surface markers, and abrogation of sphere formation and tumorigenicity *in vivo*. While telomere length can be maintained by alternative lengthening of telomere (ALT) mechanisms in 10-15% of the tumors, the activation of ALT pathways is rare in cancer and most cancers maintain telomeres through regulation of telomerase activity. (Shay & Wright, 2019) Furthermore, the telomere staining pattern observed in our cells shows no indication for an important role of ALT.

Direct promoter activation may also lead to increased TERT activity in tumor cells, and two common promoter mutations (−124 and −146 bp upstream of the TERT start site) are typically associated with increased telomerase activity. While these mutations are frequent in other cancers, they rarely occur in GI-tumors, (Vinagre *et al.*, 2013) which is well in line with our own observation that all PDAC cells we tested showed wildtype promoter sequences.

Using BIBR1532 as a small molecule telomerase inhibitor, we detected a dose-dependent proliferation arrest, which is in line with reduced cell proliferation observed in ESCs. Critical shortening of telomeres induces DNA damage response pathways, cell-cycle arrest and finally cell death. These effects are triggered by ATM and/or ATR-dependent signaling via checkpoint activation, resulting in the marking of uncapped chromosomal ends by formation of gH2AX foci and telomere dysfunction-induced foci (TIFs).(Sexton *et al*, 2014) Thus, the cellular response to dysfunctional telomeres is regulated through the same factors that control DNA damage response (Takai *et al*, 2003) and activate p53 and its downstream targets.(Brassat *et al*, 2011) We have previously demonstrated that CD133+ CSCs exhibit enhanced DNA damage repair and are particularly sensitive to inhibition of ATR-mediated DNA damage response.(Gallmeier *et al*, 2011)

Indeed, we observed that BIBR1532 treatment resulted in gH2AX foci formation and senescence in CD133+ cells. These effects were accompanied by preferential induction of apoptosis in the CSC population, emphasizing the dependency of CSCs on functional telomerase and stabilized telomeres. CSCs also appear to be more susceptible to telomerase inhibition-induced DNA damage and ultimately to the induction of apoptosis, which is especially interesting since other stem cells (ESCs and iPSCs) are particularly sensitive to DNA damage-induced apoptosis. (Liu *et al*, 2016)

Since pancreatic CSCs are highly resistant to chemotherapy, (Li *et al.*, 2007) their elimination is crucial to improve therapy efficacy and outcome for PDAC patients. We have repeatedly shown that the identification of stem cell features in CSCs, such as the activity of embryonic signaling pathways can result in novel treatment strategies to eliminate pancreatic CSCs, resulting in long-term survival of mice harboring PDAC. (Lonardo *et al.*, 2011; Mueller *et al.*, 2009) Here we demonstrate a new approach to this goal and show that pancreatic CSCs are highly dependent on telomerase activity and telomere elongation, and that loss of telomerase activity by genetic knockdown or small molecule inhibitors eliminates patient-derived pancreatic CSCs. For this study we used a series of primary cell lines generated from patient-derived xenografts. While this inevitably leads to more heterogeneity when it comes to results, it is important to note that the underlying mechanisms as well as the effects of telomerase inhibition, both by small molecules and *TERT*-KD, are reproducible in all utilized cell lines, strongly supporting the significance of the results.

The prediction of drug response *in vitro* on 2D monolayer cultures is traditionally very difficult. 3D pancreatic organoid cultures better recapitulate the situation in a tumor and are superior to 2D cultures when it comes to drug testing.(Griffith & Swartz, 2006) Therefore, we generated organoid cultures from patient-derived xenografts for drug testing and detected a dose-dependent response of organoid viability to both BIBR1532 and gemcitabine treatment. To demonstrate the efficacy of a drug regimen against CSCs, however, tumor formation (or lack thereof) *in vivo* remains the gold standard. Therefore, we used an inducible *TERT*-KD construct, allowing for treatment to commence after tumor formation and growth, replicating the therapeutic scenario in a clinical setting without the confounding off-target effects of small molecule inhibitors. Indeed, we observed significantly smaller tumors whenever telomerase function was lost, indicating a crucial role for TERT in tumor propagation.

*TERT*-KD depletes CSCs, highlighting the dependency of CSCs on intact telomerase regulation. In this respect CSCs seem resemble human ESCs, where TERT has been shown to be an essential mediator of pluripotency, cell cycle progression and differentiation.(Liu, 2017; Yang *et al*, 2008) furthermore, we observed that *TERT*-KD critically downregulates the core pluripotency factors *OCT4, KLF4, SOX2* and *NANOG,* and KLF4 is known to bind and activate the TERT promoter in human and mouse stem cells. (Hoffmeyer *et al*, 2012; Wong *et al*, 2010) However, many of the transcription factors involved in the regulation of self-renewal and pluripotency in human ESCs are tightly intertwined; KLF4 binds and upregulates the NANOG promoter, which in turn is also regulated through cooperation of SOX2 and OCT4 with KLF4.(Chan *et al*, 2009; Nakatake *et al*, 2006) Furthermore, silencing of OCT3/4 correlates with downregulation of pluripotency factors and TERT, with simultaneous activation of p53 and upregulation of its target genes p21 and PUMA.(Zhang *et al*, 2014) The subsequent p53-dependent induction of cell cycle arrest, senescence, apoptosis and the suppression of pluripotency and self-renewal in ESCs as a result of DNA damage has also been demonstrated. (Chin *et al*, 1999) Thus, a tightly coordinated regulation of stemness-associated transcription factors and pathways is essential for the preservation of “stemness”.

We therefore hypothesized that *TERT* expression is regulated by the same transcription factor signature; Using a NANOG reporter system we demonstrate that high NANOG expression correlates directly with TERT expression and vice versa, conferring increased CD133 expression and sphere formation ability to cancer cells. Using expression vectors for *OCT3/4*, *SOX2*, *KLF4* and *NANOG*, we demonstrate that while all four pluripotency factors seem to increase *TERT* expression and thus impact the regulation of telomerase activity, *SOX2* and *OCT3/4* do so most strongly. In summary, these data show a previously undiscovered positive feed-back loop between pluripotency transcription factors and TERT regulation in pancreatic CSCs.

*TERT* is furthermore regulated by epigenetic changes, such as chromatin loop structures in cells with long telomeres (Kim *et al*, 2016) or through CpG islet methylation of the *TERT* promoter. However, epigenetic changes of *TERT* by methylation remain a controversial topic: Bechter et al. demonstrated that in patients with B-CLL, *TERT* promoter hypomethylation was associated with an increased telomerase activity, (Bechter *et al*, 2002) whereas others showed that *TERT* hypermethylation increases *TERT* mRNA expression and telomerase activity in a variety of cancers (e.g. bladder, brain, colon, heart and kidney) (Guilleret *et al*, 2002). We found no differences in *TERT* promoter methylation using previously published global methylation data, suggesting that methylation is likely not a controlling factor in pancreatic CSCs.

In summary, the results presented in this study demonstrate that telomerase regulation is critical for the maintenance of stemness in CSCs. This may significantly promote our understanding of PDAC tumor biology and may eventually result in improved treatment for pancreatic cancer patients.

## Materials and Methods

### Mice, transplantation and treatment

Female 6-8 week-old athymic Nude-Foxn1nu mice were purchased from Envigo (France). For subcutaneous xenografting, single cells were resuspended in 40 μl 1:1 media:Matrigel (Invitrogen). CSC frequencies and statistical significance of the comparison were determined from extreme limiting dilution assay results using ELDA software.(Hu & Smyth, 2009) For *in vivo* treatment, animals received biweekly intraperitoneal (i.p.) injections of doxycycline (1 mg/ml). Tumor size was calculated using the formula (length x width^2^)/2. Mice were housed and maintained in laminar flow cabinets in specific pathogen-free conditions. All animal work and care were carried out following strict guidelines following German or Spanish legal regulations to ensure careful, and ethic handling of mice. The experimental protocol for animal studies was reviewed and approved by the respective government review board and institutional animal care.

### Primary pancreatic cancer cells and other cell lines

Primary human pancreatic cancer cell lines were generated from established PDX obtained under MTAs (Reference no. I409181220BSMH and I405271505PHMH) from the CNIO, Madrid, Spain, and maintained in culture as described previously. (Mueller *et al.*, 2009) U-2OS and HPDE cells have been previously described.(Lerma *et al.*, 2016) Hek293T cells were cultured in DMEM supplemented with 10% FBS, 1% Pen/Strep and 1% Glutamine, in standard conditions (37°C, 5% CO_2_). BIBR1532 (Selleckchem) was used at 80μM (3 or 7 days treatment, changing media and BIBR1532 every other day), Gemcitabine (Merck) was used at 5μM and Olaparib (Selleckchem) was used at 25μM (for 72h) for 3D cell culture unless stated otherwise.

### CSC enrichment

#### CD133 and ALDH fluorescence activated cell sorting (FACS)

For CSC enrichment primary pancreatic cancer cells were stained with CD133 (See Additional file 1 **Table 1**) or the Aldefluor™ Kit (STEMCELL™ Technologies) was performed. Cell sorting was performed using a FACSAria III (BD Bioscience).

**Table 1.** Antibodies.

|  | Application | Manufacturer |
| --- | --- | --- |
| α-hu-CD133/1-(AC133)-APC | FC | MiltenyiBiotec (Cat no. 130090826) |
| α-hu-CD133/1-(AC133)-PE | FC | MiltenyiBiotec (Cat no. 130080801) |
| α-hu-CD133/1-(AC133)-PE-VIO770 | FC | MiltenyiBiotec (Cat no. 130102358) |
| α-hu-EpCAM-APC | FC | BD Biosciences (Cat no. 347200) |
| α-γ-H2A.X (Ser139) | IF | Merck (Cat no. 05-636) |

#### Sphere culture

Spheres were cultured as described previously. (Hermann *et al.*, 2007) Briefly, cells were cultured in DMEM-F12 supplemented with B-27 (ThermoFisher Scientific) and bFGF (Novoprotein). 10,000 cells per milliliter were seeded in ultra-low attachment plates (Corning). Spheres were defined as 3-dimensional multicellular structures of ≥40μm. After 7 days, sphere formation was quantified either manually using a Leica stereomicroscope or using a CASY TT (OLS OMNI Life Science) with a 150μm capillary. Dead cells and debris were excluded from the quantification.

### Pseudorabies virus infections

The virulent Pseudorabies virus (PRV) strain PRV-NIA3 has been previously described. The parental PRV virus vBecker2 was generated by transfection of pBecker2 plasmid into Hela Tet-Off cells.(Lerma *et al.*, 2016) PRV-TER, a recombinant PRV virus in which the endogenous viral IE180 promoter was substituted with the TERT human tumor promoter, has been detailed previously.(Lerma *et al.*, 2016) For infection of HPDE and Panc185 cells in adherence, 5×10^5^ cells were seeded in 6 multi-well plates. Twenty-four hours post seeding, cells were infected with PRV-NIA3, vBecker2 or PRV-TER at a multiplicity of infection (MOI) of 0.1 TCID_50_/cell. After an absorption period of 2h at 37°C, the cells were washed with PBS to remove unabsorbed virus, and incubated for 72h at 37°C. Virus yield was determined from total lysates of infected cells on U2OS cells as previously described.(Lerma *et al.*, 2016) For infection of spheres, 7-day-old Panc185 1st generation spheres were trypsinized, cells were counted and cell suspensions of 5×10^5^ cells were seeded in 6 multi-well plates and infected with PRV-NIA3, vBecker2 or PRV-TER at a multiplicity of infection (MOI) of 0.1 TCID_50_/cell. After an absorption period of 2h at 37°C under agitation, cultures were washed with PBS to remove unabsorbed virus, and incubated for 72h at 37°C. Virus yield was determined as detailed above.

### Telomerase activity

For telomerase activity measurement, the TeloTAGGG telomerase PCR Elisa^PLUS^ kit (Roche), the telomeric repeat amplification protocol (TRAP) or qRT-PCR based TRAP were performed. Primer sequences are provided in the Supplementary Information (See Supplementary file 1 **Table 2**).

**Table 2.**
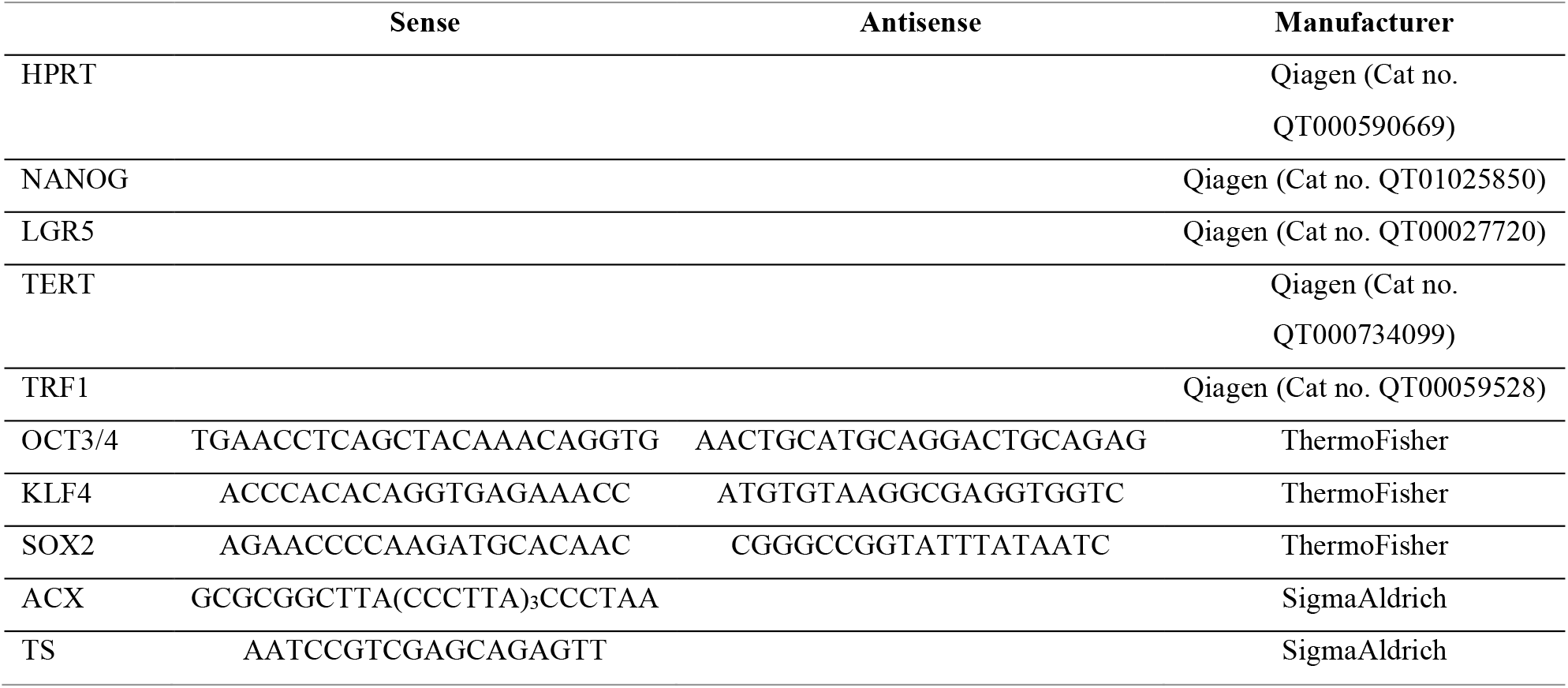
RT qPCR Primers.

### Organoid culture

Tumor pieces of patient-derived xenografts (PDXs) were digested with Accutase® solution (Merck) for 30 min at 37°C. The cells were then filtered in EASY strainer 100 μM (greiner bio-one) and cultured in Matrigel coated plates containing organoid culture medium as described in Dantes et al. 2017 with 5 % growth factor reduced Matrigel (BD, 354230). Media of the organoid culture plates was refreshed every 3-4 days. Organoids were treated with BIBR1532 and gemcitabine at 12.5, 25.0 and 50 μM for 72 h. To determine the number of metabolically active and viable organoids, the CellTiter-Glo® 3D Cell Viability Assay (Promega, G9681) was performed following the manufacturer’s instructions.

### Plasmids, infection and transfection

For RNAi-mediated gene silencing, the pGIPZ and pTRIPZ lentiviral vectors developed by Dr. Greg Hannon (Cold Spring Harbor Laboratory) and Dr. Steve Elledge (Harvard Medical School) and the corresponding shRNA constructs (GIPZ TERT shRNAs, RHS4531-EG7015 and TRIPZ TERT shRNAs, RHS4740-EG7015) were purchased from Dharmacon. The NANOG reporter lentiviral vector backbone and the sequence of the construct have been previously described. (Hotta *et al*, 2009) NANOG, OCT3/4, SOX2 and KLF4 were cloned into the PB-EF1-MCS-IRES-RFP plasmid (SBI). Transposition was performed using Super PiggyBac transposase (SBI), transfection was performed with standard Lipofectamine® 2000 (Invitrogen).

### Flow cytometry

Antibodies used in this study are listed in the Supplementary file 1 (**Table1**). For ALDH detection the Aldefluor™ kit (STEMCELL™Technologies) was used. Apoptosis was measured using Annexin V-APC (BD) and CellEvent™ Caspase-3/7 green detection reagent (invitrogen). For DNA content staining cells were stained with DAPI. Cellular senescence was measured using the SA-ß-gal kit (BioCat). Samples were analyzed by flow cytometry using a LSR II (BD Bioscience), and data were analyzed with FlowJo V10 (Ashland, Oregon). FACSorting was performed using a FACSAria III (BD).

### Q-FISH

Telomere Q-FISH was performed as previously described.(Hummel *et al*, 2015) Fluorescence intensity of the telomeres was quantified with Definiens software (Definiens, Munich, Germany).

### Immunofluorescence

Primary pancreatic cancer cells were FACSorted, seeded on cover-slips in 6-well dishes, incubated at 37°C for 12h, washed with PBS and fixed with 2% PFA for 20min at room temperature. Coverslips were washed 3 times with PBS and incubated at room temperature for 15min in 0.7% triton in PBS. 1h blocking with 5% milk powder in 0.1% TBS-T at room temperature was followed by incubation with the respective antibodies (1:500) in a humidified chamber at 4°C overnight. Coverslips were then washed 3x with 0.1% TBS-T. Secondary antibodies (1:1,000) were incubated at room temperature in a humidified chamber for 2h, washed and mounted with ProLong™Gold Antifade mounting reagent with DAPI (ThermoFisher). Images were captured using a Leica TCS SP8-HCS confocal microscope. Antibodies are listed in the Supplementary file (**Table 1**).

### RNA isolation and qRT-PCR

Total RNA was prepared using RNeasy kits with on-column genomic DNA digestion following the manufacturer’s instructions (Qiagen). First strand cDNA was prepared using QuantiTect Reverse Transcription kit (Qiagen). Reactions were performed with QuantiFastSybr Green PCR Kit (Qiagen) using a QuantStudio 3 machine (Applied Biosystems). Results were analyzed using the 2^−ddCt^ method and calculated as relative to *HPRT* expression. Reactions were carried out from at least three independent experiments. Primer sequences are provided in the Supplementary file (**Table 2**).

### MTT assay

1,000 cells were seeded in a 96 well plate and incubated for 24h at 37°C. Cells were incubated for 72h with the indicated concentrations of the respective drugs and further incubated for 3h with 5 mg/ml MTT (Merck). Finally, DMSO (Roth) was added and the optical density was measured at 560nm using an Infinite Pro 200 plate reader (Tecan, Switzerland).

### Statistical Analysis

Results for continuous variables are presented as means ± SEM. Unless stated otherwise, treatment groups were compared with the one-sided Mann-Whitney-U test, tumor growth dynamics were compared by calculating areas under the curve. Chi-squared tests were used to compare CSC frequencies. Contingency tables were compared using Fisher’s exact test. P values <0.05 were considered statistically significant. Statistical analyses were performed using GraphPad Prism 5.0 (San Diego, CA).

### Data availability

This study includes no data deposited in external repositories. All data generated and analyzed in this study are included within the article and are available from the corresponding author.

## Acknowledgments

We are indebted to Andrea Wißmann and Kristina Diepold for excellent technical and experimental support. This report includes data that were generated as part of the doctoral thesis of K.W. at Ulm University. Technical illustrations in this manuscript were used from Servier Medical Art under a creative commons attribution 3.0 unported license.

## Author contribution

K.W., F.B., B.S.Jr., and P.C.H. conceived the study and designed the experiments. K.W., T.D., K.T., L.L., L.A.S., F.A., V.U., N.A., M.E., A.L., and B.S.Jr. performed the experiments and analyzed the data. M.V.F., F.B., and T.B. designed and performed telomere length analyses. S.M., and C.G., provided technical support for TERT-knockdown constructs. K.W., E.R.A., P.O.F., T.S., A.K., F.B., B.S.Jr., and P.C.H. contributed to the study design, as well as writing and revising the manuscript. B.S.Jr., and P.C.H. acquired funding for the project. All authors contributed to the review and editing of the manuscript.

## The Paper Explained Problem

Despite intensive research pancreatic ductal adenocarcinoma (PDAC) still has a poor prognosis and the therapeutic options are limited. Cancer stem cells (CSCs) are essential for PDAC propagation, for its metastatic spread and chemoresistance. To date the precise mechanism, which preserves the stemness characteristics of CSCs and regulates the unlimited self-renewal potential of these cells is unclear.

## Results

We discovered that telomerase activity and telomere elongation are essential for the maintenance of the stemness characteristics in pancreatic CSCs. Stemness in pancreatic CSCs is regulated via a feedback loop between stemness factors (NANOG, OCT3/4, SOX2, KLF4) and telomerase activity. The inhibition of telomerase via small molecule inhibitor treatment or genetic knock-down diminished CSCs in human primary pancreatic cancer cells by inducing DNA damage and apoptosis.

## Impact

Our study demonstrates that telomerase regulation is critical for stemness maintenance in pancreatic CSCs and examines the effects of telomerase inhibition as a potential treatment option of pancreatic cancer. This significantly promotes our understanding of PDAC tumor biology and CSC regulation and may result in improved treatment for pancreatic cancer patients.

## Figure legends

**Supplementary Figure 1.**
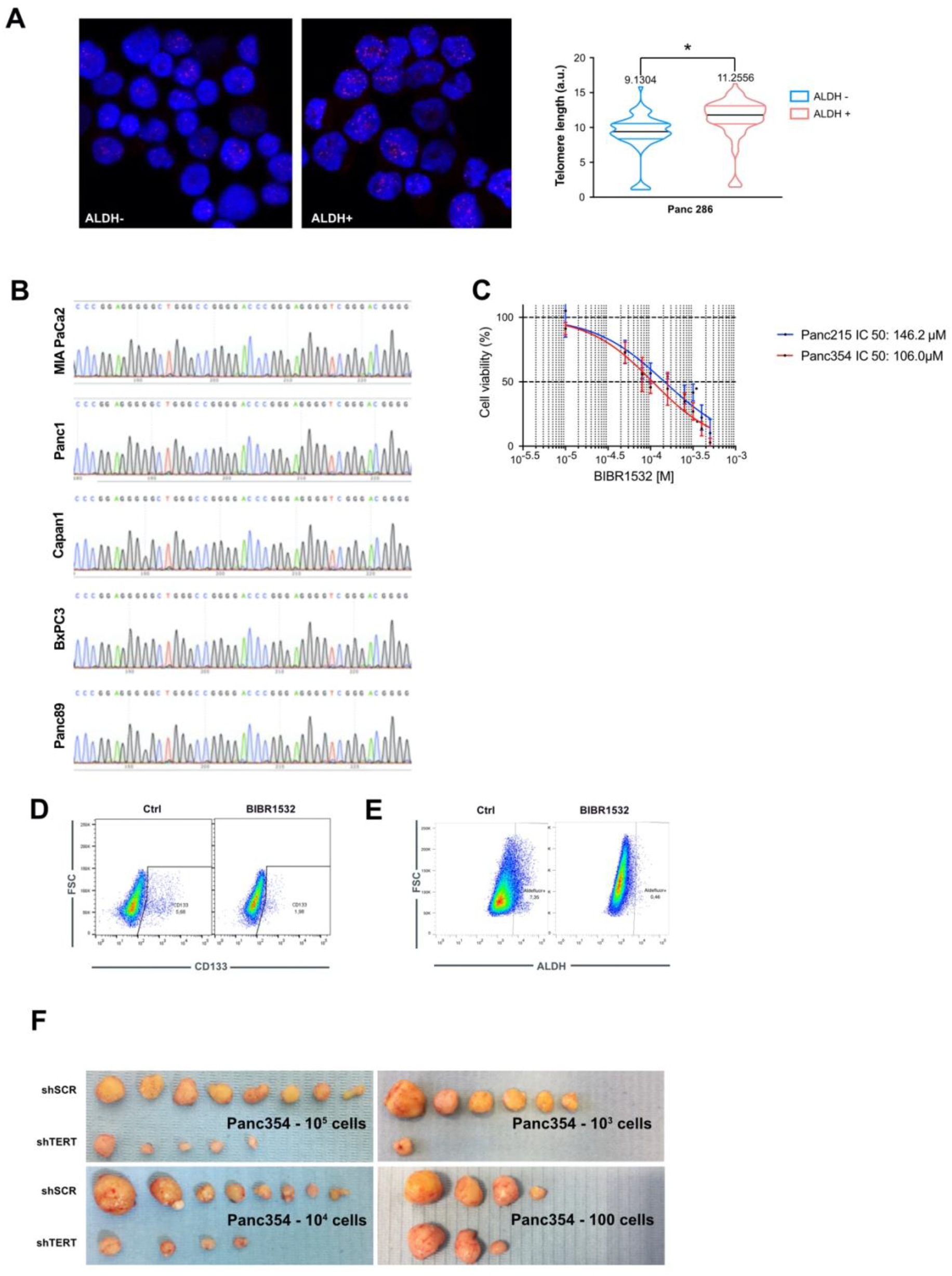
Additional Data. A Representative pictures and Q-FISH telomere length analysis in Aldefluor positive and negative cells. The mean is depicted in numbers and as black line, >150 measurements per group. B TERT promoter mutation analysis of C250T and C228T sites in five established pancreatic cancer cell lines. C MTT assay to determine the respective IC_50_ of BIBR1532 in the utilized primary pancreatic cancer cells. D, E Representative cytometry illustrations of CD133 (D) and Aldefluor (E) positive cells after vehicle (Ctrl) or BIBR1532 treatment. F Representative pictures of explanted subcutaneous tumors from Panc354 ELDA assays. In (A) data are represented as mean ± SEM. *P ≤ 0.05 (Mann-Whitney-U test).

